# Identifying outlier loci in admixed and in continuous populations using ancestral population differentiation statistics

**DOI:** 10.1101/054585

**Authors:** Helena Martins, Kevin Caye, Keurcien Luu, Michael G.B. Blum, Olivier François

## Abstract

Finding genetic signatures of local adaptation is of great interest for many population genetic studies. Common approaches to sorting selective loci from their genomic background focus on the extreme values of the fixation index, *F*_ST_, across loci. However, the computation of the fixation index becomes challenging when the population is genetically continuous, when predefining subpopulations is a difficult task, and in the presence of admixed individuals in the sample. In this study, we present a new method to identify loci under selection based on an extension of the *F*_ST_ statistic to samples with admixed individuals. In our approach, *F*_ST_ values are computed from the ancestry coefficients obtained with ancestry estimation programs. More specifically, we used factor models to estimate *F*_ST_, and we compared our neutrality tests with those derived from a principal component analysis approach. The performances of the tests were illustrated using simulated data, and by re-analyzing genomic data from European lines of the plant species *Arabidopsis thaliana* and human genomic data from the population reference sample, POPRES.

## 1 Introduction

Natural selection, the process by which organisms that are best adapted to their environment have an increased contribution of genetic variants to future generations, is the driving force of evolution (Darwin, 1859). Identifying genomic regions that have been the targets of natural selection is one of the most important challenge in modern population genetics (Vitti *et al*., 2013). To this aim, examining the variation in allele frequencies between populations is a frequently applied strategy (Cavalli-Sforza, 1966). More specifically, by sampling a large number of single nucleotide polymorphisms (SNPs) throughout the genome, loci that have been affected by diversifying selection can be identified as outliers in the upper tail of the empirical distribution of *F*_ST_ (Lewontin & Krakauer, 1973; Beaumont & Nichols, 1996; Akey *et al*., 2002; Weir *et al*., 2005). For selectively neutral SNPs, *F*_ST_ is determined by migration and genetic drift, which affect all SNPs across the genome in a similar way. In contrast, natural selection has locus-specific effects that can cause deviations in *F*_ST_ values at selected SNPs and at linked loci.

Outlier tests based on the empirical distribution of *F*_ST_ across the genome requires that the sample is subdivided into *K* subsamples, each of them corresponding to a distinct genetic group. For outlier tests, defining subpopulations may be a difficult task, especially when the background levels of *F*_ST_ are weak and when populations are genetically homogeneous (Waples & Gaggiotti, 2006). For example, Europe is genetically homogeneous for human genomes, and it is characterized by gradual variation in allele frequencies from the south to the north of the continent (Lao *et al*., 2008), in which genetic proximity mimics geographic proximity (Novembre *et al*., 2008). Studying evolution in the field, most ecological studies use individual-based sampling along geographic transects without using prior knowledge of populations (Manel *et al*., 2003; Schoville *et al*., 2012). For example, the 1001 genomes project for the plant species *Arabidopsis thaliana* used a strategy in which individual ecotypes were sampled with a large geographic coverage of the native and naturalized ranges (Horton *et al*., 2012; Weigel & Mott, 2009). One last difficulty with *F*_ST_ tests arises from the presence of individuals with multiple ancestries (admixture), for which the genome exhibits a mosaic of fragments originating from different ancestral populations (Long, 1991). The admixture phenomenon is ubiquitous over sexually reproducing organisms (Pritchard *et al*., 2000). Admixture is pervasive in humans because migratory movements have brought together peoples from different origins (Cavalli-Sforza *et al*., 1994). Striking examples include the genetic history of African American and Mestizo populations, for which the contributions of European, Native American, and African populations had been studied extensively (Bryc *et al*., 2010; Tang *et al*., 2007).

Most of the concerns raised by definitions of subpopulations are commonly answered by the application of clustering or ancestry estimation approaches such as `structure` or principal component analysis (PCA) (Pritchard *et al*., 2000; Patterson *et al*., 2006). These approaches rely on the framework of factor models, where a factor matrix, the *Q*-matrix for `structure` and the score matrix for PCA, is used to define individual ancestry coefficients, or to assign individuals to their most probable ancestral genetic group (Engelhardt & Stephens, 2010). To account for geographic patterns of genetic variation produced by complex demographic histories, spatially explicit versions of the `structure` algorithm can include models for which individuals at nearby locations tend to be more closely related than individuals from distant locations (François & Durand, 2010).

In this study, we propose new tests to identify outlier loci in admixed and in continuous populations by extending the definition of *F*_ST_ to this framework (Long, 1991). Our tests are based on the computation of ancestry coefficient and ancestral allele frequency, Q and F, matrices obtained from ancestry estimation programs. We develop a theory for the derivation of this new *F*_ST_ statistic, defining it as the proportion of genetic diversity due to allele frequency differences among populations in a model with admixed individuals. Then we compute our new statistic using the outputs of two ancestry estimation programs: `snmf` which is used as fast and accurate version of the `structure` algorithm, and `tess3` a fast ancestry estimation program using genetic and geographic data (Frichot *et al*., 2014; Caye *et al*., 2016). Using simulated data sets and SNPs from human and plants, we compared the results of genome scans obtained with our new *F*_ST_ statistic with the results of PCA-based methods (Hao *et al*., 2016; Duforet-Frebourg *et al*., 2016; Chen *et al*., 2016; Galinsky *et al*., 2016; Luu *et al*., 2016).

## 2 *F*-statistics for populations with admixed individuals

In this section, we extend the definition of *F*_ST_ to populations containing admixed individuals, and for which no subpopulations can be defined a priori. We consider SNP data for *n* individuals genotyped at *L* loci. The data for each individual, *i*, and for each locus, *ℓ*, are recorded into a genotypic matrix *Y*. The matrix entries, *y_iℓ_*, correspond to the number of derived or reference alleles at each locus. For diploid organisms, *y_iℓ_* is an integer value 0, 1 or 2.

### A new definition of *F*_ST_

Suppose that a population contains admixed individuals, and the source populations are unknown. Assume that individual ancestry coefficients, *Q*, and ancestral population frequencies, *F*, are estimated from the genotypic matrix *Y* by using an ancestry estimation algorithm such as `structure` (Pritchard *et al*., 2000). Consider a particular locus, *ℓ*, and let *f_k_* be the reference allele frequency in ancestral population *k* at that locus. We set

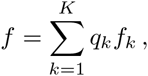

where *q_k_* is the average value of the population *k* ancestry coefficient over all individuals in the sample, and the ancestral allele frequencies are obtained from the *F* matrix. Our formula for *F*_ST_ is

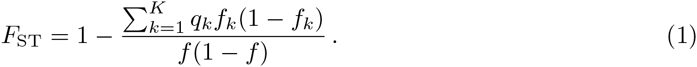

The above definition of *F*_ST_ for admixed populations is obviously related to the original definition of Wright’s fixation index. Assuming *K* predefined subpopulations, Wright’s definition of *F*_ST_ writes as follows (Wright, 1951)

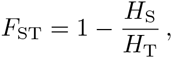

where 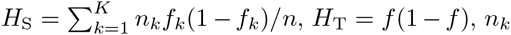 is the sample size, *f_k_* is the allele frequency in subpopulation *k*, and *f* is the allele frequency in the total population. For admixed samples, the estimates of the sample sizes, *n_k_*, are obtained by setting *n_k_* = *nq_k_*, and the sampled allele frequencies are replaced by their ancestral allele frequencies. The interpretation of the new *F*_ST_ statistic is thus similar to the interpretation of Wright’s fixation index. The main distinction is its application to ‘idealized’ ancestral populations inferred by `structure` or a similar algorithm. For recently admixed populations, our new statistic represents a measure of population differentiation due to population structure prior to the admixture event. Mathematically rigorous arguments for this analogy will be given in a subsequent paragraph.

### Admixture estimates

While many algorithms can compute the *Q* and *F* matrices, our application of the above definition will focus on ancestry estimates obtained by nonnegative matrix factorization algorithms (Frichot *et al*., 2014). Frichot *et al*. (2014)’s algorithm runs faster than the Monte-Carlo algorithm implemented in `structure` and than the optimization methods implemented in `faststructure` or `admixture` (Alexander *et al*., 2009; Raj *et al*., 2014). Estimates of *Q* and *F* matrices obtained by the `snmf` algorithm can replace those obtained by the program `structure` advantageously for large SNP data sets (Wollstein & Lao, 2015).

The `snmf` algorithm estimates the *F* matrix as follows. Assume that the sampled genotype frequencies can be modelled by a mixture of ancestral genotype frequencies

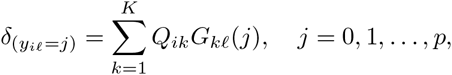

where *y_iℓ_* is the genotype of individual *i* at locus *ℓ*, the *Q_ik_* are the ancestry coefficients for individual *i* in population *k*, the *G_kℓ_*(*j*) are the ancestral genotype frequencies in population *k*, and *p* is the ploidy of the studied organism (*δ* is the Kronecker delta symbol indicating the absence/presence of genotype *j*). For diploids (*p* = 2), the relationship between ancestral allele and genotype frequencies can be written as follows

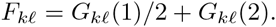

The above equation implies that the sampled allele frequencies, *x_iℓ_*, satisfy the following equation

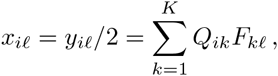

which makes the estimates consistent with the definition of *F*_ST_.

### Population differentiation tests

The regression framework explained in the next paragraph leads to a direct approximation of the distribution of *F*_ST_ under the null-hypothesis of a random mating population (Sokal & Rohlf, 2012). In this framework, we define the squared *z*-scores as follows

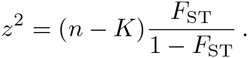

Assuming random mating at the population level, we have

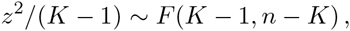

where *F*(*K* − 1,*n* − *K*) is the Fisher distribution with *K* − 1 and *n* − *K* degrees of freedom. In addition, we assume that the sample size is large enough to approximate the distribution of squared *z*-scores as a chi-squared distribution with *K* − 1 degrees of freedom.

A naive application of this theory would lead to an increased number of false positive tests due to population structure. In genome scans, we adopt an empirical null-hypothesis testing approach which recalibrates the null-hypothesis. The principle of test calibration is to evaluate the levels of population differentiation that are expected at selectively neutral SNPs, and modify the null-hypothesis accordingly (François *et al*., 2016). Following GWAS approaches, this can be achieved after computing the genomic inflation factor, defined by the median of the squared *z*-scores divided by the median of a chi-squared distribution with *K* − 1 degrees of freedom (genomic control, Devlin & Roeder (1999)).

### Software

The methods described in this section were implemented in the R package LEA (Frichot & François, 2015). A short tutorial on how to compute the *F*_ST_ statistic and implement the tests is available at http://goo.gl/OsRhLQ.

### Mathematical theory

A classical definition for the fixation index, *F*_ST_, corresponds to the proportion of the genetic variation (or variance) in sampled allele frequency that can be explained by population structure

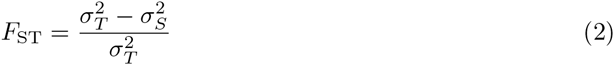

where, in the analysis of variance terminology, 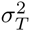 is the total variance and 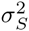 is the error variance (Weir, 1996). This definition of *F*_ST_, which uses a linear regression framework, can be extended to models with admixed individuals in a straightforward manner. Suppose that a population contains admixed individuals, and assume we have computed estimates of the *Q* and *F* matrices. For diploid organisms, a genotype is the sum of two parental gametes, taking the values 0 or 1. In an admixture model, the two gametes can be sampled either from the same or from distinct ancestral populations. The admixture model assumes that individuals mate randomly at the moment of the admixture event. Omitting the locus subscript *ℓ*, a statistical model for an admixed genotype at a given locus can be written as follows

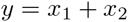

where *x*_1_ and *x*_2_ are independent Bernoulli random variables modelling the parental gametes. The conditional distribution of *x*_1_ (resp. *x*_2_) is such that prob(*x*_1_ = 1|Anc_1_ = *k*) = *f_k_* where *f_k_* is the allele frequency in ancestral population *k*, Anc is an integer value between 1 and *K* representing the hidden ancestry of each gamete. The sampled allele frequency is defined as *x* = *y*/2 (*x* taking its values in 0, 1/2, 1). Thus the expected value of the random variable *x* is given by the following formula

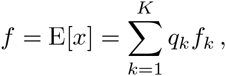

where *q_k_* = prob(Anc = *k*). The total variance of *x* satisfies

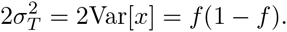

Using the *Q* and *F* matrices, *q_k_* can be estimated as the average value of the ancestry coefficients over all individuals in the sample, and the ancestral allele frequencies can be estimated as *f_k_* = *F_k_*.

To compute the error variance, 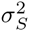, we consider that the two gametes originate from the same ancestral population. Assuming Hardy-Weinberg equilibrium in the ancestral populations, the error variance can be computed as follows

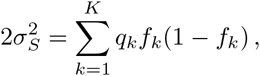

and the use of equation (2) for *F*_ST_ concludes the proof of equation (1).

## 3 Simulation experiments and data sets

### Simple simulation models

In a first series of simulations, we created replicate data sets close to the underlying assumptions of population differentiation tests (Lewontin & Krakauer, 1973; Beaumont & Nichols, 1996). While relying on simplified assumptions, those easily reproducible simulations have the advantage of providing a clear ‘proof-of-concept’ framework which connects our new statistic to the classical theory. Admixed genotypes from a unique continuous population were obtained from two ancestral gene pools. In this continuous population, individual ancestry varied gradually along a longitudinal axis. The samples contained 200 individuals genotyped at 10,000 unlinked SNPs. Ancestral polymorphisms were simulated based on Wright’s two-island models. Two values for the proportion of loci under selection were considered (5% and 10%). To generate genetic variation at outlier loci, we assumed that adaptive SNPs had migration rates smaller than the migration rate at selectively neutral SNPs. In this model, adaptive loci experienced reduced levels of ancestral gene flow compared to the genomic background (Bazin *et al*., 2010). The effective migration rate at a neutral SNP was equal to one of the four values 4*Nm* = 20,15,10, 5. The effective migration rate at an adaptive SNP was equal to one of the four values 4*Nm_s_* = 0.1, 0.25, 0.5,1. A total number of 32 different data sets were generated by using the computer program ms (Hudson, 2002).

The model for admixture was based on a gradual variation of ancestry proportions across geographic space (Durand *et al*., 2009). Geographic coordinates (*x_i_*,*y_i_*) were created for each individual from Gaussian distributions centered around two centroids put at distance 2 on a longitudinal axis (standard deviation [SD] = 1). As it happens in a secondary contact zone, we assumed that the ancestry proportions had a sigmoidal shape across space (Barton & Hewitt, 1985),

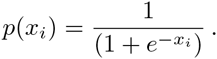

For each individual, we assumed that each allele originated in the first ancestral population with probability *p*(*x_i_*) and in the second ancestral population with probability 1 − *p*(*x_i_*) (Durand *et al*., 2009).

### Complex simulation models

To evaluate the power of tests in realistic landscape simulations, we used six publicly available data sets previously described by Lotterhos & Whitlock (2015). In those scenarios, the demographic history of a fictive species corresponded to nonequilibrium isolation by distance due to expansion from two refugia. The simulations mimicked a natural population whose ranges have expanded since the last glacial maximum, potentially resulting in secondary contact (Hewitt, 2000). The study area was modelled as a square with 360 x 360 demes. Migration was determined by a dispersal kernel with standard deviation *σ* = 1.3 demes, and the carrying capacity per deme was 124. The data sets consisted of 9900 neutral loci and 100 selected loci. Twenty unrelated individuals were sampled from thirty randomly chosen demes. For each replicate data set, a selective landscape was randomly generated based on spherical models described as ‘weak clines’ (details in Lotterhos & Whitlock (2015)). All selected loci adapted to this landscape.

### Computer programs

We performed genome scans for selection using three factor methods: `snmf` (Frichot *et al*., 2014), `tess3` (Caye *et al*., 2016), `pcadapt` (Luu *et al*., 2016; Duforet-Frebourg *et al*., 2016). A fourth method used the standard *F*_ST_ statistic where subpopulations were obtained from the assignment of individuals to their most likely genetic cluster. Like for `snmf`, the `tess3` estimates of the *Q* and *G* matrices are based on matrix factorization techniques. The main difference between the two programs is that tess3 computes ancestry estimates by incorporating information on individual geographic coordinates in its algorithm whereas the `snmf` algorithm is closer to `structure` (Caye *et al*., 2016). The default values of the two programs were implemented for all their internal parameters. Each run of the two programs was replicated five times, and the run with the lowest cross-entropy value was selected for computing *F*_ST_ statistics according to formula (1). We compared the results of `snmf` and `tess3` with the results of the program `pcadapt` (Luu *et al*., 2016). The test statistic of the latest version of `pcadapt` is the Manhanalobis distance relative to the *z*-scores obtained after regressing the SNP frequencies on the *K* − 1 principal components. As for `snmf` and for `tess3`, test calibration in `pcadapt` was based on the computation of the genomic inflation factor. For genome scans based on the *F*_ST_ statistic where subpopulations are obtained from the assignment of individuals to their most likely genetic cluster, we used a chi-squared distribution with *K* — 1 of freedom after recalibration of the null-hypothesis using genomic control. Before applying the methods to the simulated data sets, the SNPs were filtered out and only the loci with minor allele frequency greater than 5% were retained for analysis.

### Real data sets

To provide an application of our method to natural populations, we reanalyzed data from the model plant organism *Arabidopsis thaliana*. This annual plant is native to Europe and central Asia, and within its native range, it goes through numerous climatic conditions and selective pressures (Mitchell-Olds & Schmitt, 2006). We analyzed genomic data from 120 European lines of *A. thaliana* genotyped for 216k SNPs, with a density of one SNP per 500 bp (Atwell *et al*., 2010). To reduce the sensitivity of methods to an unbalanced sampling design, fourteen ecotypes from Northern Scandinavia were not included in our analysis. Those fourteen ecotypes represented a small divergent genetic cluster in the original data set. In addition to the plant data, we analyzed human genetic data for 1,385 European individuals genotyped at 447k SNPs (Nelson *et al*., 2008).

### Candidate lists

After recalibration of the null-hypothesis using genomic inflation factors, histograms of test significance values were checked for displaying their correct shape. Then, False Discovered Rate (FDR) control algorithms were applied to significance values using the Storey and Tibishirani algorithm (Storey & Tibshirani, 2003). For simulated data, lists of outlier loci were obtained for an expected FDR value of 10%. The same nominal level was applied for the analysis of the human data set. For *A. thaliana*, an expected FDR value of 1% was applied, and a consensus list of loci was obtained by including all peak values present in Manhattan plots for `snmf` and `tess3`.

## 4 Results

### Simple simulation models

We evaluated the performances of genome scans using tests based on `snmf, tess3, pcadapt`, and *F*_ST_, in the presence of admixed individuals. For `snmf` and for `tess3`, we used *K* = 2 ancestral populations. This value of *K* corresponded to the minimum of the cross-entropy criterion when *K* was varied in the range 1 to 6, and it also corresponded to the true number of ancestral populations in the simulations. We used `pcadapt` with its first principal component. Considering expected FDR values between 0.01 and 0.2, we computed observed FDR values for the lists of outlier loci produced by each test. The observed FDR values remained generally below their expected values (Figure 1 for data sets with 5% of loci under selection, Figure S1 for data sets with 10% of loci under selection). These observations confirmed that the use of genomic inflation factors leads to overly conservative tests (Francois *et al*., 2016). Since similar levels of observed FDR values were observed across the 4 tests, we did not implement other calibration methods than genomic control.

**Figure 1.**
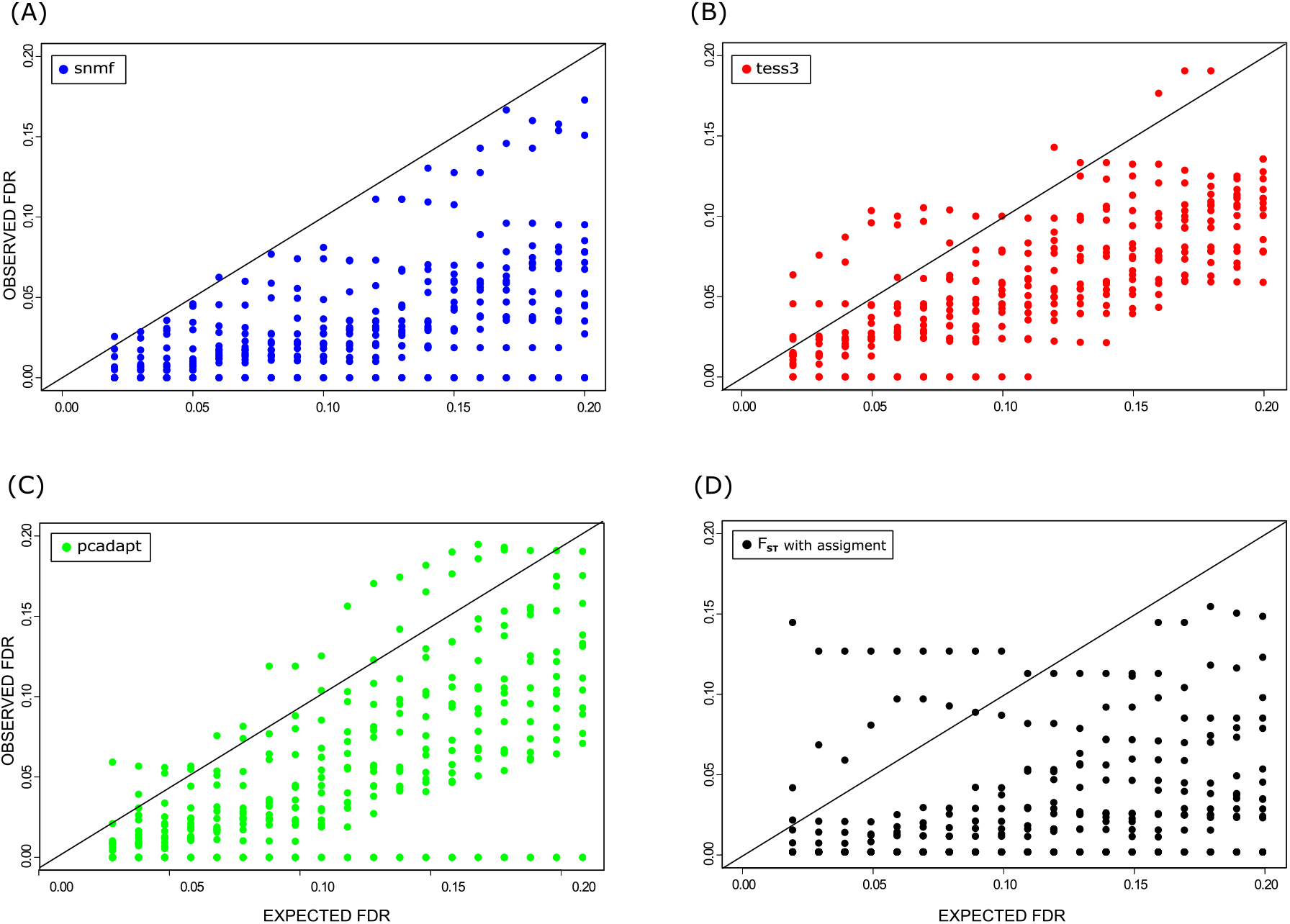
FDR for simulations of admixed populations. Simulation of ancestral populations based on 2-island models with various levels of population differentiation and selection. Sixteen data sets contained 5% of truly selected loci. Observed false discovery rates for an expected level of FDR equal to 0.1. (A) *F*_ST_ tests based on `snmf` *Q* and *F* matrices, (B) *F*_ST_ tests based on `tess3` *Q* and *F* matrices, (C) Luu et al.’s (2016) `pcadapt` statistic, (D) Standard *F*_ST_ test based on assignment of individuals to their most likely genetic cluster.

Next, we evaluated the sensitivity (power) of the four tests in each simulation scenario. Our experiments confirmed that the use of approaches that estimate ancestry coefficients is appropriate when no subpopulation can be predefined (Figure 2A for ancestry coefficient estimates). As we expected from the simulation process, the tests had higher power when the relative levels of selection intensity were higher. For 4*Nm* = 5 and 4*Nm_s_* = 0.1,0.25, 0.5, and 1, the power of tests for `snmf, tess3, pcadapt` was close to 27% for data sets with 5% of outliers (Figure 2B, expected FDR equal to 10%). The *F*_ST_ test based on assignment of individuals to their most likely cluster failed to detect outlier loci (power value equal to 0%). For 4*Nm* = 10, the power of the tests ranged between 40% and 45% for `snmf, tess3, pcadapt`, and it was equal to 26% for the *F*_ST_ test (Figure 2B). For 4*mN* ≥ 15, corresponding to the highest selection rates, the power was approximately equal to 50% for all methods considered. The relatively low power values confirmed that the tests were conservative, and truly-adaptive loci were difficult to detect. To provide an upper bound on the power of outlier tests in the context of admixed populations, we applied an *F*_ST_ test to the samples obtained prior to admixture, estimating allele frequencies from their true ancestral populations. For 4*Nm* = 5 and 10, the power of the tests for `snmf, tess3, pcadapt` was similar to the power obtained when we applied outlier tests to the data before admixture (Figure 2B). The results for data sets with 10% of selected loci were similar to those obtained with 5% of selected loci (Figure S2).

**Figure 2.**
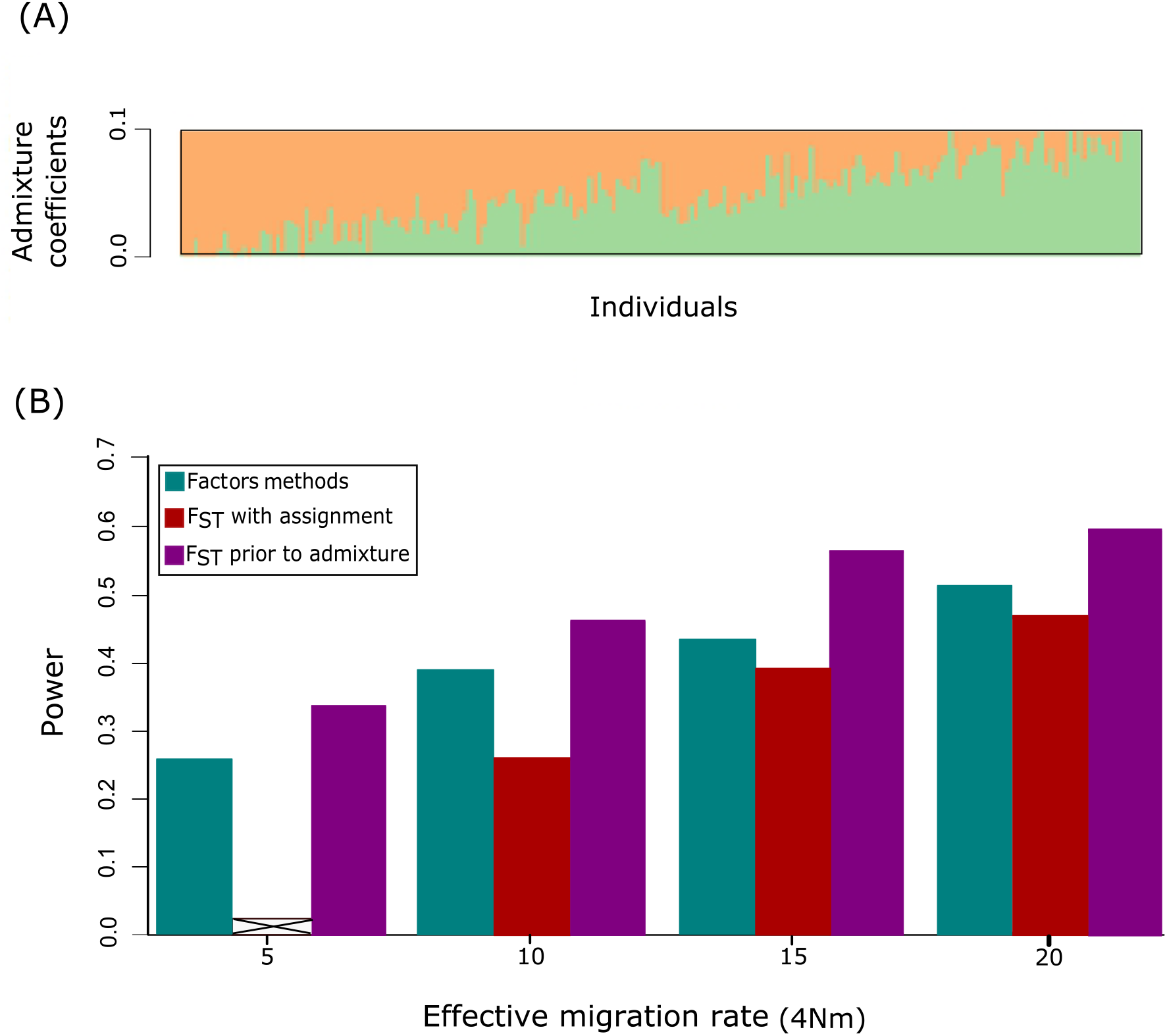
Power in simulations of admixed populations. Simulations of ancestral populations based on 2-island models with various levels of selection and background of levels of population differentiation (4*Nm*). Sixteen data sets contained 5% of truly selected loci. (A) Individual ancestry coefficients estimated from neutral loci using `snmf` with *K* = 2. (B) Power estimates for tests based on factor methods (grouping `snmf, tess3` and `pcadapt`), for *F*_ST_ tests in which individuals were assigned to their most likely cluster, and for *F*_ST_ tests prior to admixture. Power values were computed by considering an expected FDR value equal to 0.1. For 4*Nm* = 5 (relatively weak selection intensity), the *F*_ST_ test based on assignment failed to detect outlier loci.

### Complex simulation models

We compared the power of factor methods to the power of tests based on assignment of individuals to their most likely cluster in realistic landscape simulations (Lotterhos & Whitlock, 2015). As a consequence of isolation by distance, the cross-entropy curve for `snmf` decreased with the value of the number of clusters, but the curve did not exhibit a minumum. A plateau reached at *K* = 6 indicated that this value of *K* could be the best choice for modelling the mixed levels of ancestry in the data (Figure 3A). In agreement with this result, `pcadapt` consistently found 5 axes of variation in the data. For values of *K* = 4 – 7 and for an expected level of FDR of 10%, the power of tests based on factor methods ranged between 0.82 and 0.87 (Figure 3B). Although SNP rankings were not different for `pcadapt`, the `pcadapt` tests were less conservative than the tests based on the default values of `snmf` (values not reported). Classical tests that assigned individuals to their most likely cluster had power ranging between 0.44 and 0.48. The power values for classical *F*_ST_ tests were substantially lower than those obtained with the new tests.

**Figure 3.**
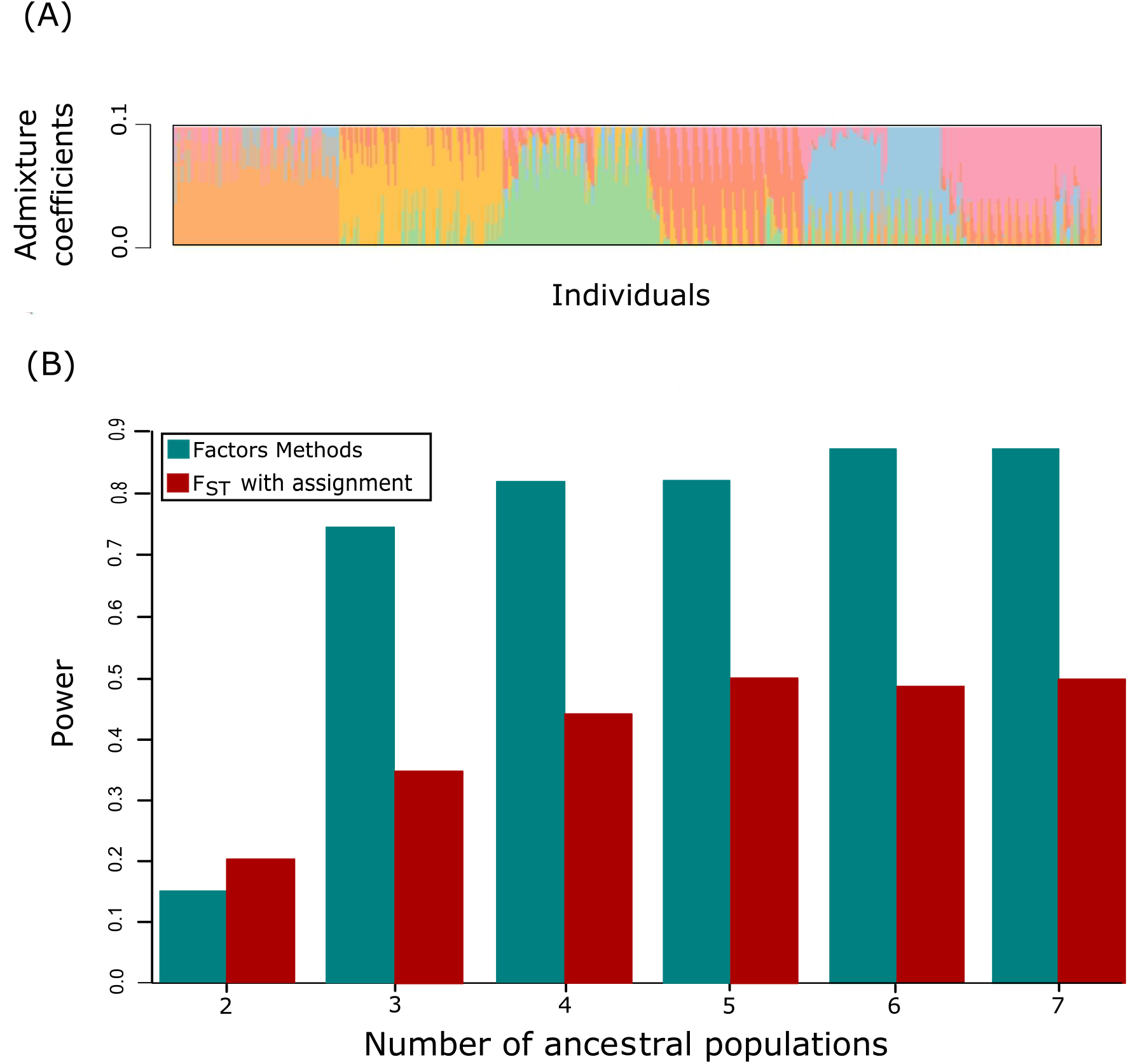
Power in simulations of range expansions. (A) Individual ancestry coefficients estimated using `snmf` with *K* = 6 ancestral populations. (B) Power estimates for tests based on factor methods and for *F*_ST_ tests in which individuals were assigned to their most likely cluster. Power values were computed by considering an expected FDR value equal to 0.1. Factor methods included `snmf` and `pcadapt`.

### Arabidopsis data

We applied `snmf, tess3` and `pcadapt` to perform genome scans for selection in 120 European lines of *Arabidopsis thaliana* (216k SNPs). Each ecotype was collected from a unique geographic location, and there were no predefined populations. To study adaptation at the continental scale, a small number ecotypes from Northern Scandinavia, which were grouped by clustering programs, were removed from the original data set of Atwell *et al*. (2010). For `snmf` and `tess3`, the cross-entropy criterion indicated that there are two main clusters in Europe, and that finer substructure could be detected as a result of historical isolation-by-distance processes. For *K* = 2, the western cluster grouped all lines from the British Isles, France and Iberia and the eastern cluster grouped all lines from Germany, and from Central and Eastern Europe (Figure 4). For implementing genome scans for selection, we used two clusters in `snmf` and `tess3`, and one principal component in `pcadapt`. The genomic inflation factor was equal to λ = 11.5 for the test based on `snmf`, and it was equal to λ = 13.1 for the test based on `tess3`. The interpretation of these two values is that the background level of population differentiation that was tested in `snmf` and `tess3` is around 0.09 (François et al. 2016). For the three methods, the Manhattan plots exhibited peaks at the same chromosome positions (Figure 5). For an expected FDR level equal to 1%, the Storey and Tibshirani algorithm resulted in a list of 572 chromosome positions for the `snmf` tests and 882 for the `tess3` tests. Figure S3 displays a Manhattan plot for the plant genome showing the main outlier loci detected by our genome scans for selection for *K* = 2. Unlike for simulated data, the tests based on PCA were more conservative than the tests based on genetic clusters. Generally, the differences between test significance values among methods could be attributed to the estimation of the genomic inflation factor and test calibration issues rather than to strong differences in SNP ranking. The results of genome scans for selection were also investigated for values of *K* greater than 2. The higher values of *K* revealed additional candidate genomic regions that were consistently discovered by the three factor methods (Figures S4–S6).

**Figure 4.**
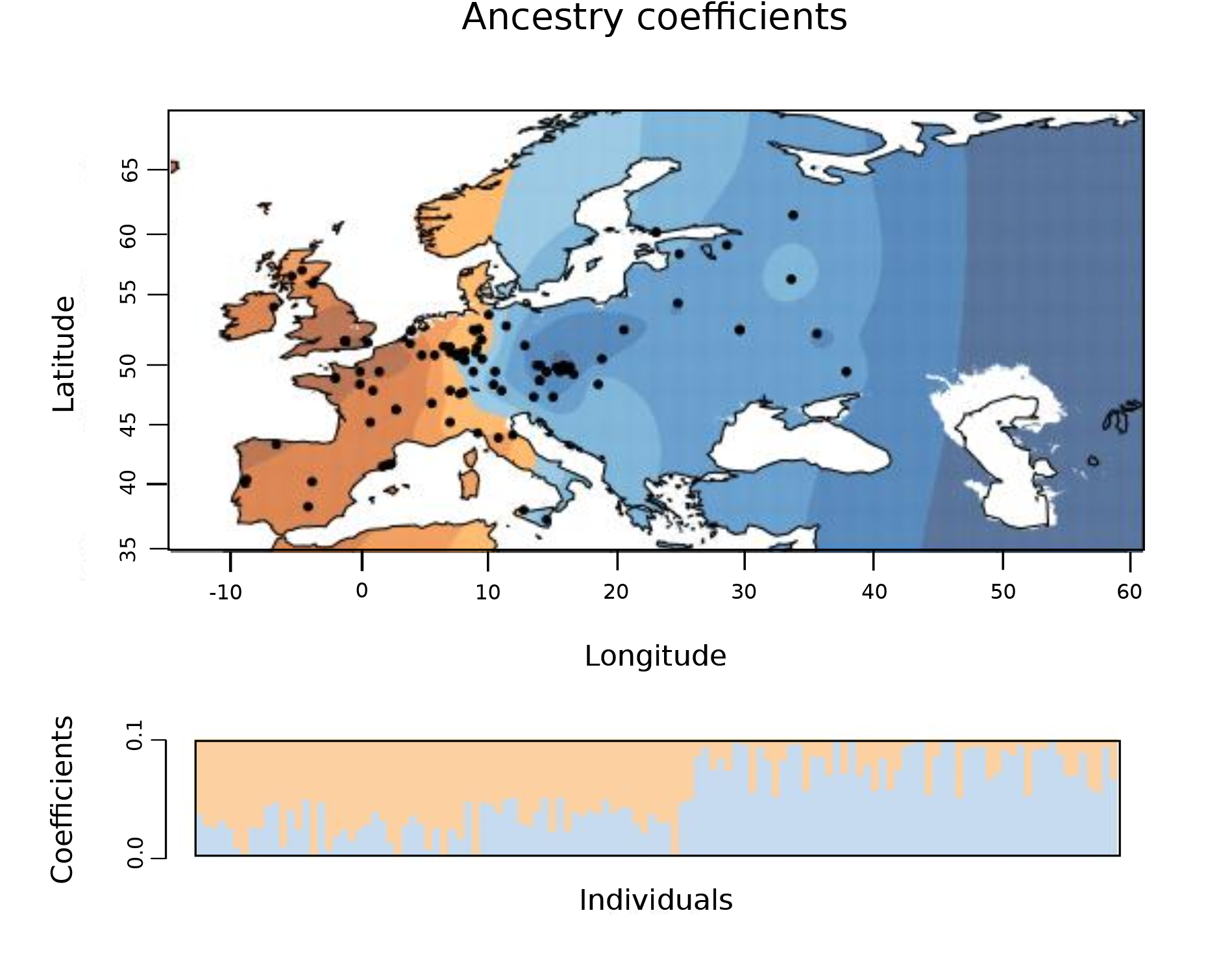
Ancestry coefficients for *Arabidopsis thaliana*. Coefficients estimated using `snmf` with *K* = 2 ancestral populations interpolated on a geographic map of Europe.

**Figure 5.**
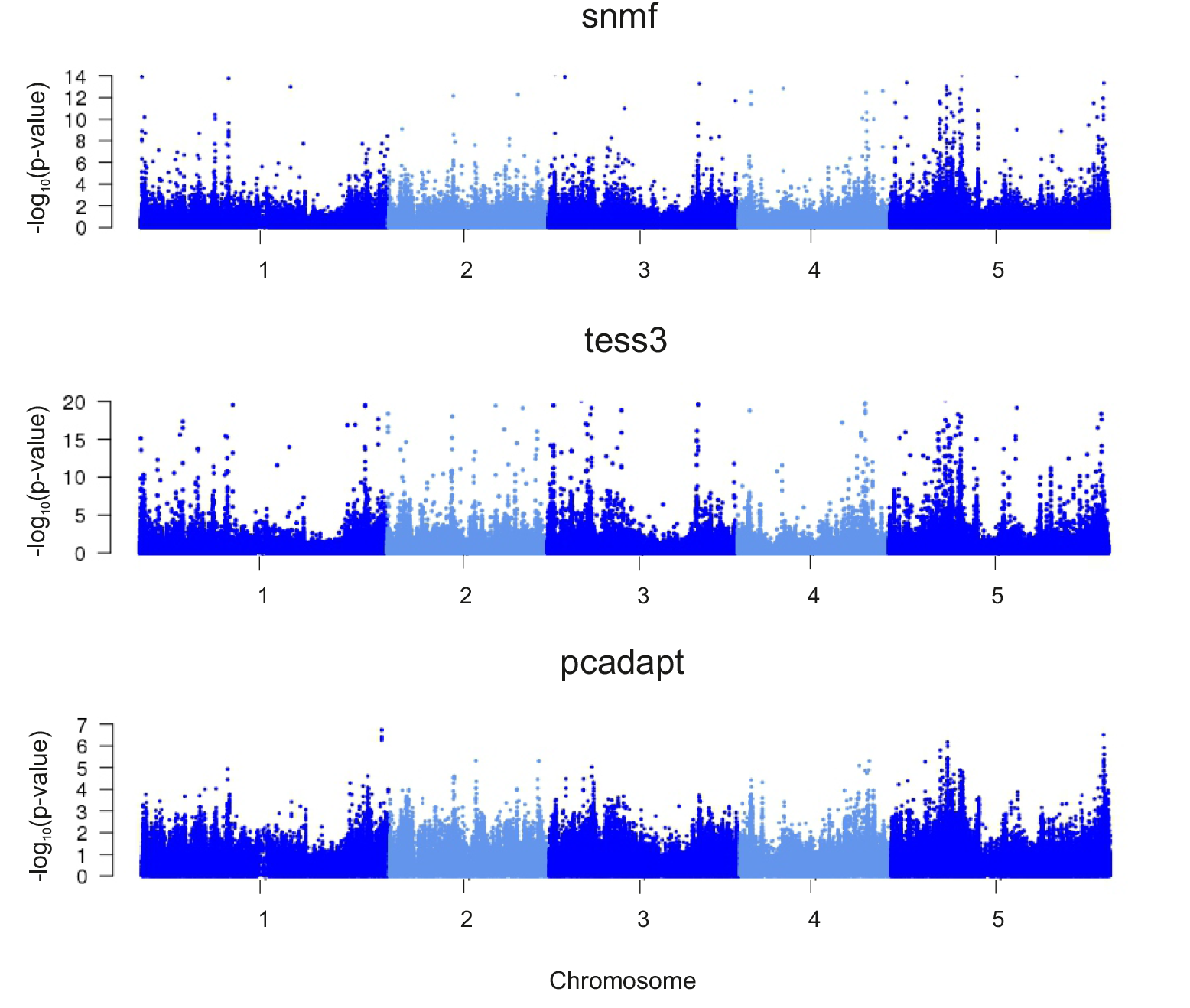
Manhattan plots of minus log10(p-values) for *A. thaliana*. Tests using (A) `snmf`, (B) `tess3` and (C) `pcadapt`. The tests based on `pcadapt` were more conservative than the tests based on the other methods.

Table 1 reports a list of 33 candidate SNPs for European *A. thaliana* lines in the 10% top hits, based on the peaks detected by the factor methods. For chromosome 1, the list contains SNPs in the gene AT1G80680 involved in resistance against bacterial pathogens. For chromosome 2, the list contains SNPs in the gene AT2G18440 (AtGUT15), which can be used by plants as a sensor to interrelated temperatures, and which has a role for controlling growth and development in response to a shifting environment (Lu *et al*., 2005). For chromosome 3, the list contains SNPs in the gene AT3G11920 involved in cell redox homeostasis. Fine control of cellular redox homeostasis is important for integrated regulation of plant defense and acclimatory responses (Muhlenbock *et al*., 2007). For chromosome 4, we found SNPs in the gene AT4G31180 (IBI1) involved in defense response to fungi. The most important list of candidate SNPs was found in the fifth chromosome. For example, the list of outlier SNPs contained SNPs in the gene AT5G02820, involved in endoreduplication, that might contribute to the adaptation to adverse environmental factors, allowing the maintenance of growth under stress conditions (Chevalier *et al*., 2011), in the genes AT5G18620, AT5G18630 and AT5G20620 (UBIQUITIN 4) involved in response to temperature stress (Kim & Kang, 2005), and in the gene AT5G20610 which is involved in response to blue light (DeBlasio *et al*., 2005). Several additional candidates were found with values of *K* greater than two for the `snmf` tests. For *K* = 3 and *K* = 4 those additional outlier regions included one SNP in the flowering locus FRIGIDA and four SNPs in COP1-interacting protein 4.1 on chromosome 4 (Horton *et al*. (2012), Figure S6). For the tests with *K* = 4, outlier regions included two SNPs in the FLOWERING LOCUS C (FLC) and five SNPs in the DELAY OF GEMINATION 1 (DOG1) locus (Horton *et al*. (2012), Figure S6).

**Table 1.**
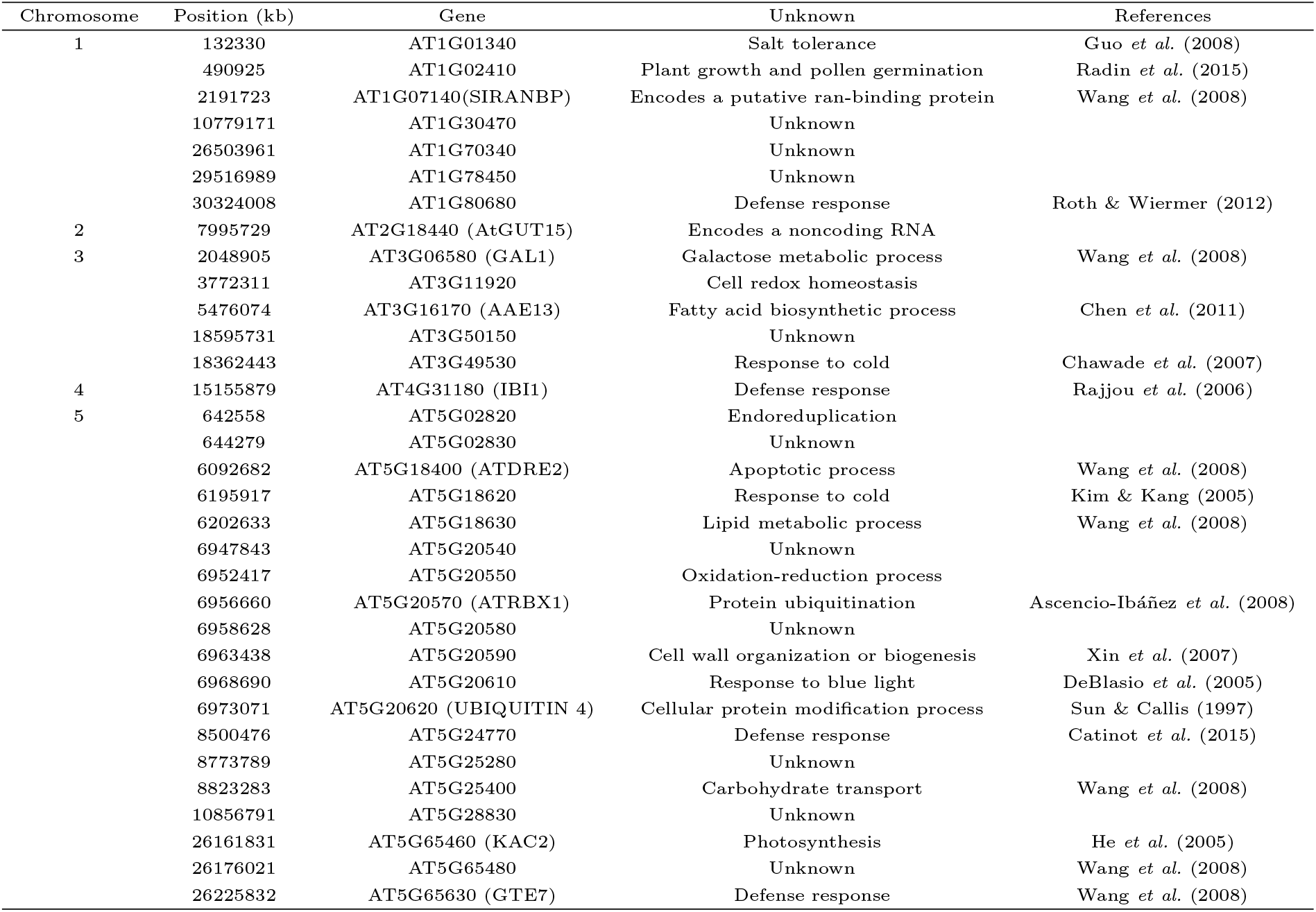
List of 33 candidate SNPs for European ecotypes of *A. thaliana*. The list was based on the list of *p*-values obtained by using an expected FDR of 1% for `snmf` and `tess3` tests.

### Human data

We applied the `snmf` and `pcadapt` tests to 1,385 European individuals from the POPRES data set (447k SNPs in 22 chromosomes). We used *K* = 2 ancestral populations in `snmf` and one principal component for PCA. For `snmf`, the genomic inflation factor was equal to λ = 9.0, indicating a background level of population differentiation around 0.006 between northern and southern European populations (Figure 6). For an expected FDR equal to 10%, we found 205 outlier loci using `snmf` tests, and 165 outlier loci with `pcadapt`. For chromosome 2, the most important signal of selection was found at the lactase persistence gene *(LCT)* (Bersaglieri *et al*., 2004). For chromosome 4, 5 SNPs were found at the *ADH1C* locus that is involved in alcohol metabolism (Han *et al*., 2007), close to the *ADH1B* locus reported by Galinsky *et al*. (2016). For chromosome 6, a signal of selection corresponding to the human leukocyte antigen *(HLA)* region was identified. For chromosome 15, there was an outlier SNP in the *HERC2* gene, which modulates human pigmentation (Visser *et al*. (2012), Figure 6).

**Figure 6.**
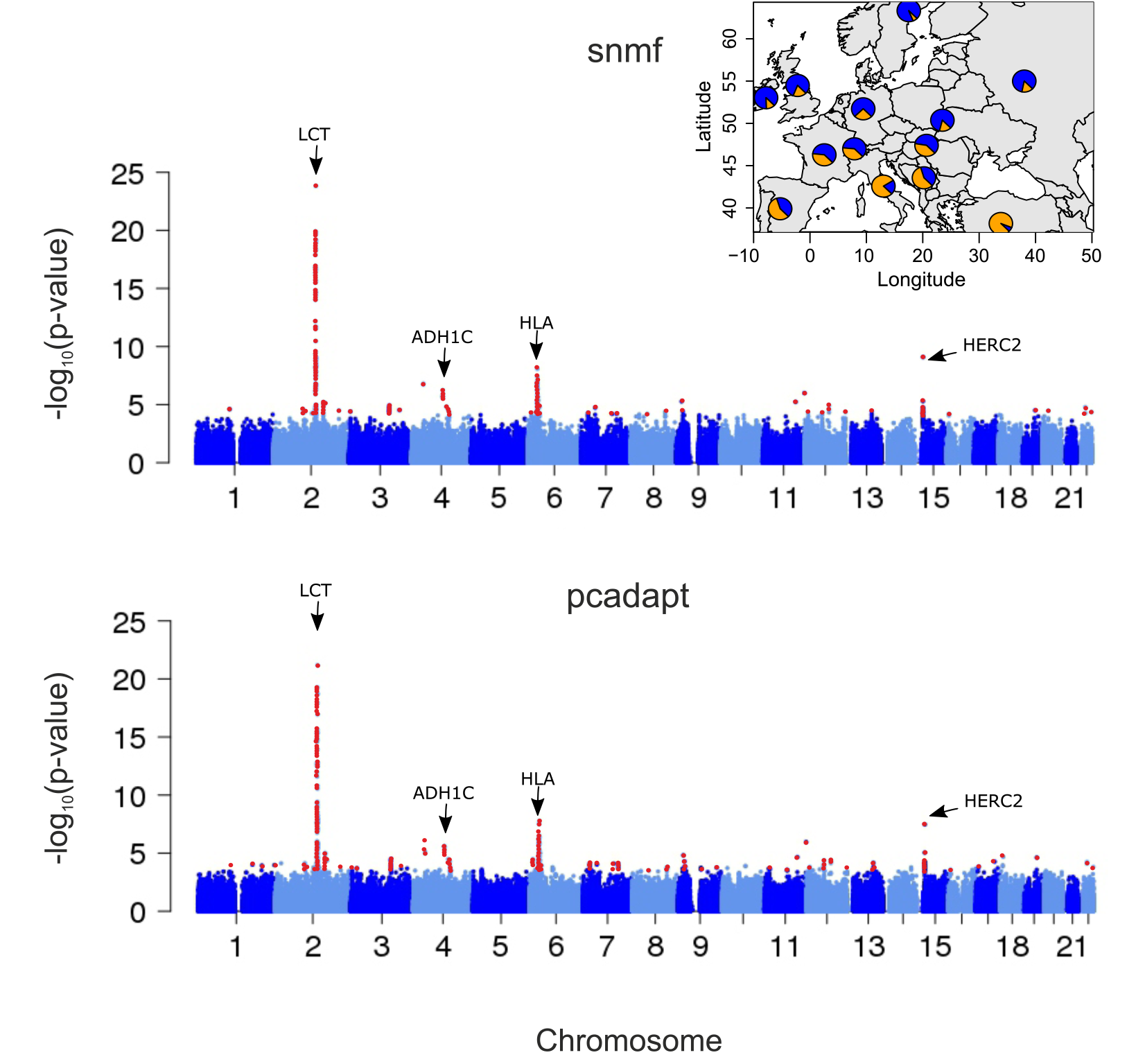
Manhattan plots of minus log10(p-values) for Europeans (POPRES data set). Tests using (A) `snmf` and (B) `pcadapt`. Candidate loci detected by genome scans for selection are colored in red for an expected FDR level of 10%. The inserted figure displays population structure estimated with `snmf` with *K* = 2 populations.

## 5 Discussion

When no subpopulation can be defined a priori, analysis of population structure commonly relies on the computation of the *Q* (and *F*) ancestry matrix obtained through the application of the program structure or one of its improved versions (Pritchard *et al*., 2000; Tang *et al*., 2005; Chen *et al*., 2007; Alexander *et al*., 2009; Raj *et al*., 2014; Frichot *et al*., 2014; Caye *et al*., 2016). In this context, we proposed a definition of *F*_ST_ based on the *Q* and *F* matrices, and we used this new statistic to screen genomes for signatures of diversifying selection. By modelling admixed genotypes, our definition of *F*_ST_ was inspired by an analysis of variance approach for the genotypic data (Weir & Cockerham, 1984; Holsinger & Weir, 2009).

The estimator for *F*_ST_ presented here is related to the estimator proposed by Long (1991) for population data. Long’s estimator was obtained from the variance of allele frequencies with respect to their expectations based on an admixture model, that enables estimating the effect of genetic drift and the effective size of the hybrid population. In order to obtain Long’s estimate, multiple locus samples are required from the hybrid population and from all contributing parental populations. For the method proposed in our manuscript, information on ancestral genetic diversity is evaluated with less prior assumptions by the application of ancestry estimation programs.

Ancestry coefficients computed by structure or similar programs are conceptual abstractions that do not always reflect demographic history correctly (Kalinowski, 2011; Puechmaille, 2016; Falush *et al*., 2016). Assuming that a large number of SNPs are genotyped across multiple populations, the calibration of statistical tests of neutrality do not require assumptions about population demographic history. Our simulations of admixed populations provided evidence that the tests based on this new statistic had an increased power compared to tests in which we assigned individuals to their most probable cluster. Interestingly, the power of those tests was only slightly lower than standard *F*_ST_ tests based on the truly ancestral allele frequencies. Going beyond simplified simulation scenarios, we evaluated the power of our tests in range expansion scenarios with complex patterns of isolation by distance. In those scenarios, genetic correlation among samples inflates the variance of population differentiation statistics (Bierne *et al*., 2013). We observed that inflation factor corrections reduced this problem when using numbers of clusters (K) greater than 2. Although a ‘true’ value for *K* did not exist, we found that the power of our tests was optimal for *K* estimated from a PCA or by cross-validation using our factor model. In this case, the ancestry coefficients disagreed with the known demographic history (simulated organisms expanded from two refugia), but the gain in performance in favor of the new tests was even higher than in the simple proof-of-concept simulations tailored to the new method.

Our reanalysis of European *A. thaliana* genetic polymorphisms provided a clear example of the usefulness of our *F*_ST_ statistic to detect targets of natural selection in plants. European ecotypes of *Arabidopsis thaliana* are continuously distributed across the continent, with population structure influenced by historical isolation-by-distance processes (Atwell *et al*., 2010; Hancock *et al*., 2011; Francois *et al*., 2008). The application of our *F*_ST_ statistic to the SNP data suggested several new candidate loci involved in resistance against pathogens, in growth and development in response to a shifting environment, in the regulation of plant defense and acclimatory responses, in the adaptation to adverse environmental factors, in allowing the maintenance of growth under stress conditions, in response to temperature stress or response to light.

An alternative approach to investigating population structure without predefined populations is by using principal component analysis (Patterson *et al*., 2006). Statistics extending the definition of *F*_ST_ were also proposed for PCA (Hao *et al*., 2016; Duforet-Frebourg *et al*., 2016; Galinsky *et al*., 2016; Chen *et al*., 2016). The performances of PCA statistics and our new *F*_ST_ statistic were highly similar. The small differences observed for the two tests could be ascribed to the chi-squared distribution approximation and to the estimation of inflation factors to calibrate the null-hypothesis. The idea of detecting signatures of selection in an admixed population has a considerable history and has been explored since the early seventies (Blumberg & Hesser, 1971; Adams & Ward, 1973; Tang *et al*., 2007). The connection between our definition of *F*_ST_ and previous works shows that the methods studied in this study, including PCA or ancestry programs, are extensions of classical methods of detection of selection using admixed populations (Long, 1991). Our results allow us to hypothesize that the age of selection detected by PCA and by our new method is similar. Thus it is likely that the selective sweeps detected by PCA and *F*_ST_ methods correspond to ancient selective sweeps already differentiating in ancestral populations. A comparison of our results for Europeans from the POPRES data sets and the genome-wide patterns of selection in 230 ancient Eurasians provides additional evidence that the signals detected by our *F*_ST_ were already present in the populations that were ancestral to modern Europeans (Mathieson *et al*., 2015).

While only minor differences between the ranking of *p*-values with 4 methods were observed, the results might be still sensitive to the algorithm used to estimating the ancestry matrices. Wollstein & Lao (2015) performed an extensive comparison of 3 recently proposed ancestry estimation methods, `admixture, faststructure, snmf` (Alexander & Lange, 2011; Raj *et al*., 2014; Frichot *et al*., 2014), and they concluded that the accuracy of the methods could differ in some simulation scenarios. In practice, it would be wise to apply several methods and to combine their results by using a meta-analysis approach as demonstrated in François *et al*. (2016).

## Data Accessibility

Simulated data are available from Lotterhos KE, Whitlock MC (2015) Data from: The relative power of genome scans to detect local adaptation depends on sampling design and statistical method. Dryad Digital Repository: http://dx.doi.org/10.5061/dryad.mh67v.

The Atwell *et al*. (2010) data are publicly available from https://github.com/Gregor-Mendel-Institute/atpolydb.

The POPRES data were obtained from dbGaP (accession number phs000145.v1.p1).

## Aknowlegments

We are grateful to three anonymous reviewers for their time and efforts in evaluating our manuscript. Helena Martins acknowledges support from the ‘Ciências sem Fronteiras’ scholarship program from the Brazilian government. This work has been partially supported by the LabEx PERSYVAL-Lab (ANR-11-LABX-0025-01) funded by the French program Investissement d’Avenir, and by the ANR AGRHUM project (ANR-14-CE02-0003-01). OF acknowledges support from Grenoble INP, and from the ‘Agence Nationale de la Recherche’ (project AFRICROP ANR-13-BSV7-0017).

H.M., K.C. and K.L. performed the analyses. H.M. and O.F. drafted the manuscript. O.F. and M.G.B.B designed the study. All authors read and approved the final version of the manuscript.

---

**Figure S1.**
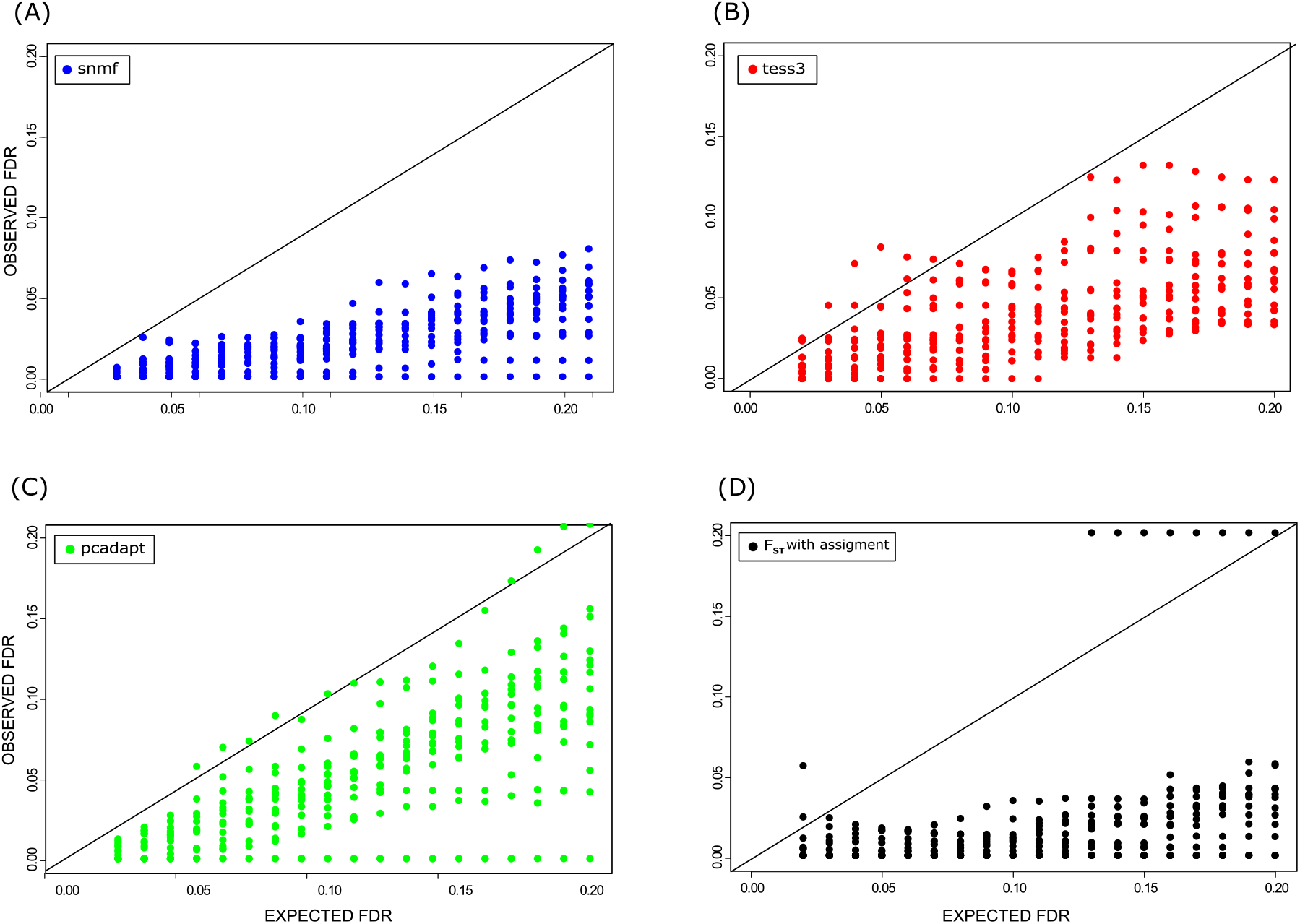
FDR for simulations of admixed populations (10% of outliers). Simulation of ancestral populations based on 2-island models with various levels of population differentiation and selection. Sixteen data sets contained 10% of truly selected loci. Observed false discovery rates for an expected level of FDR equal to 0.1. (A) *F*_ST_ tests based on `snmf` *Q* and *F* matrices, (B) *F*_ST_ tests based on `tess3` *Q* and *F* matrices, (C) Luu et al.’s (2016) `pcadapt` statistic, (D) Standard *F*_ST_ test based on assignment of individuals to their most likely genetic cluster.

**Figure S2.**
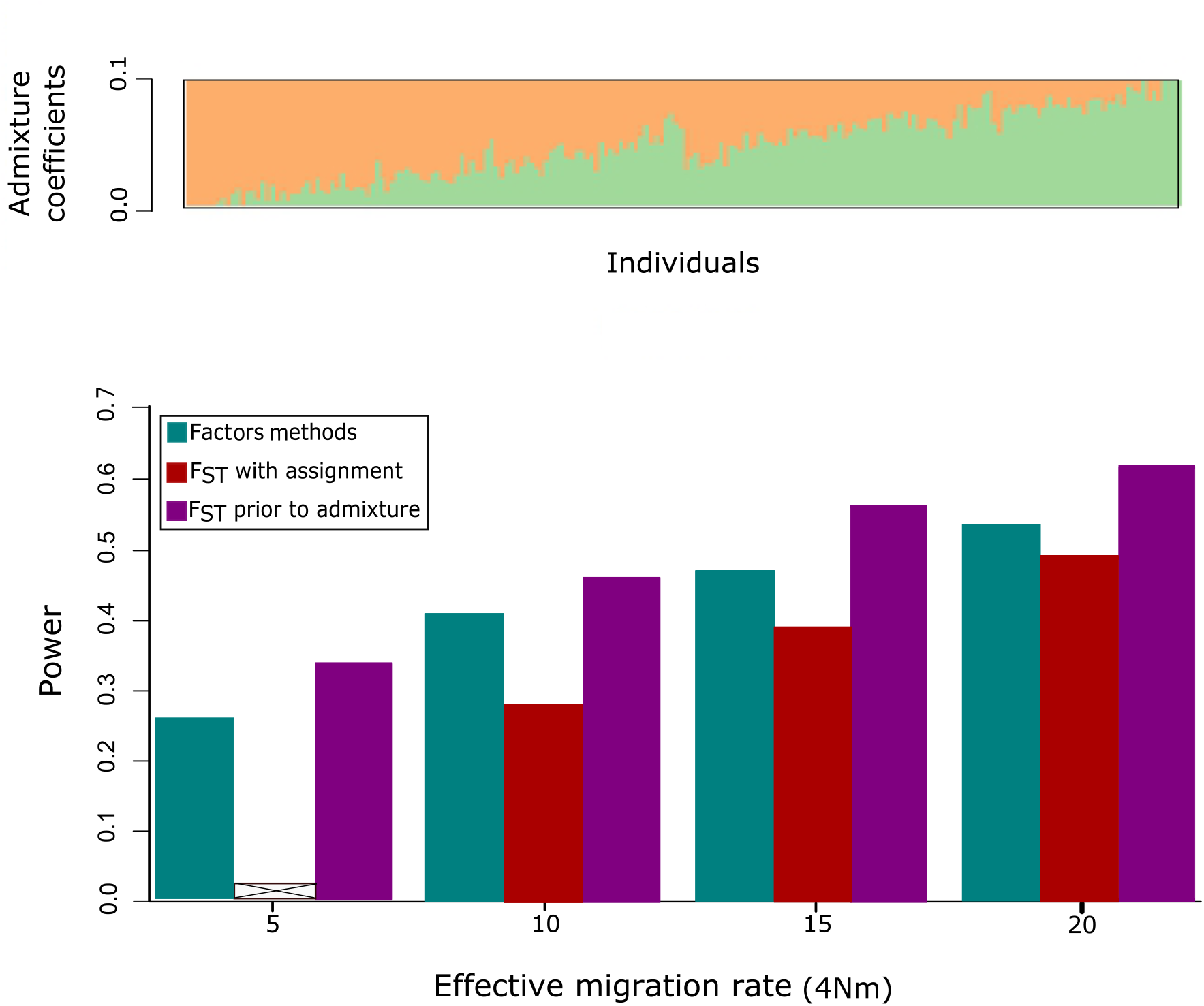
Power in simulations of admixed populations (10% of outliers). Simulations of ancestral populations based on 2-island models with various levels of selection and background of levels of population differentiation (4*Nm*). Sixteen data sets contained 10% of truly selected loci. (A) Individual ancestry coefficients estimated from neutral loci using `snmf` with *K* = 2. (B) Power estimates for tests based on factor methods (grouping `snmf, tess3` and `pcadapt`), for *F*_ST_ tests in which individuals were assigned to their most likely cluster, and for *F*_ST_ tests prior to admixture. Power values were computed by considering an expected FDR value equal to 0.1. For 4*Nm* = 5 (weak selection intensity), the *F*_ST_ test based on assignment failed to detect outlier loci.

**Figure S3.**
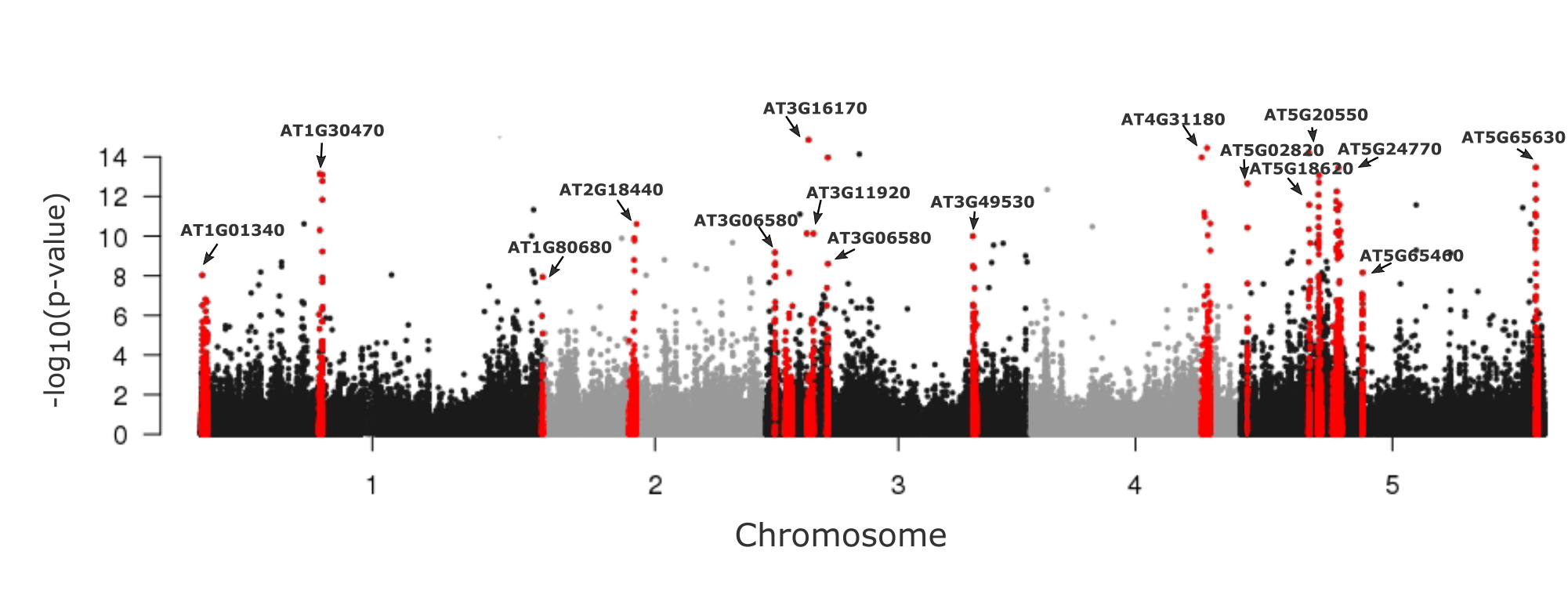
Manhattan plot of minus log10(p-values) for *A. thaliana*. The candidate regions are colored in red. Those regions correspond to an expected FDR level of 1% for `snmf` and `tess3` having more than 5 SNPs in each region.

**Figure S4.**
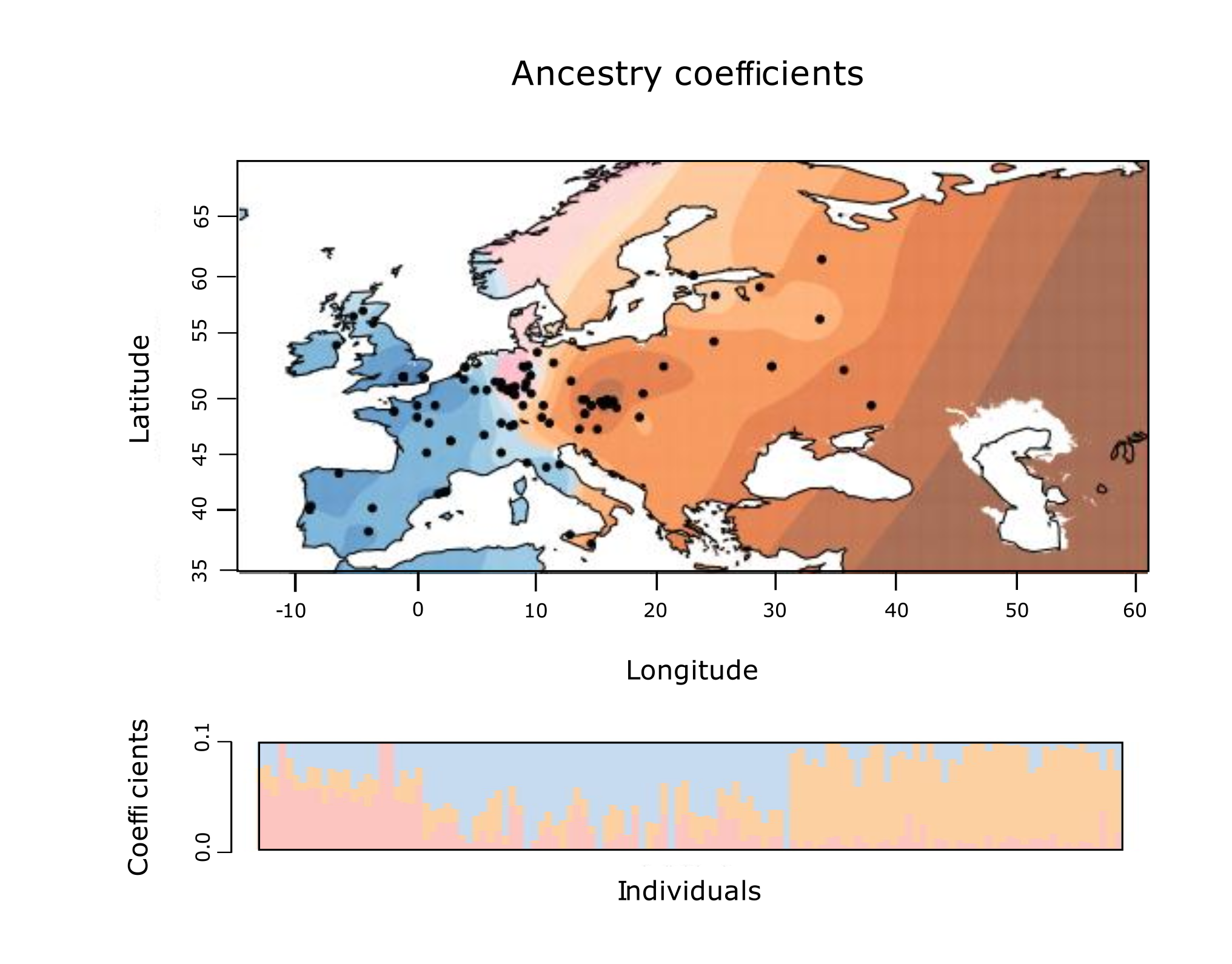
Geographic map of ancestry coefficients for *Arabidopsis thaliana* using snmf with *K* = 3 ancestral populations.

**Figure S5.**
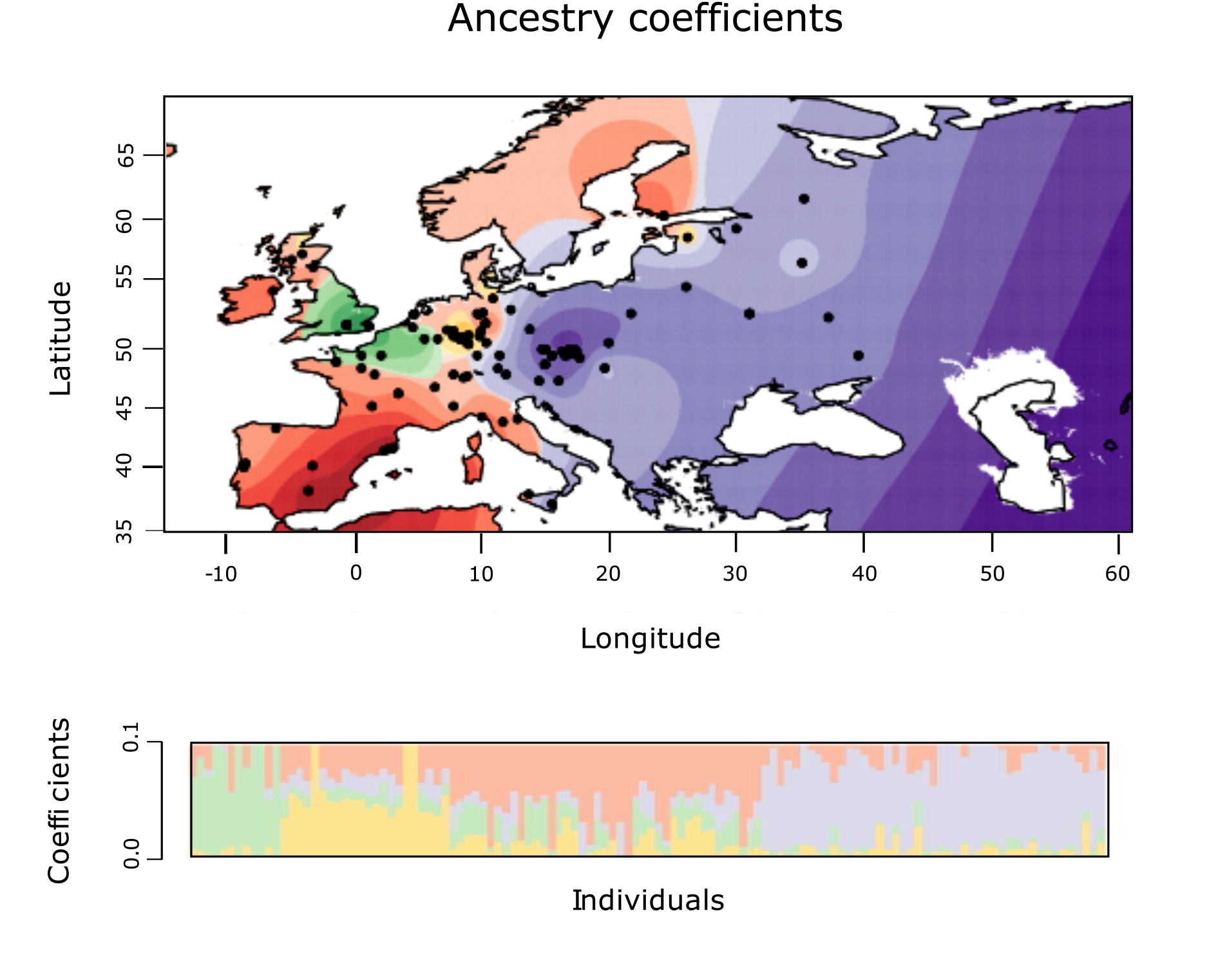
Geographic map of ancestry coefficients for *Arabidopsis thaliana* using snmf with *K* = 4 ancestral populations.

**Figure S6.**
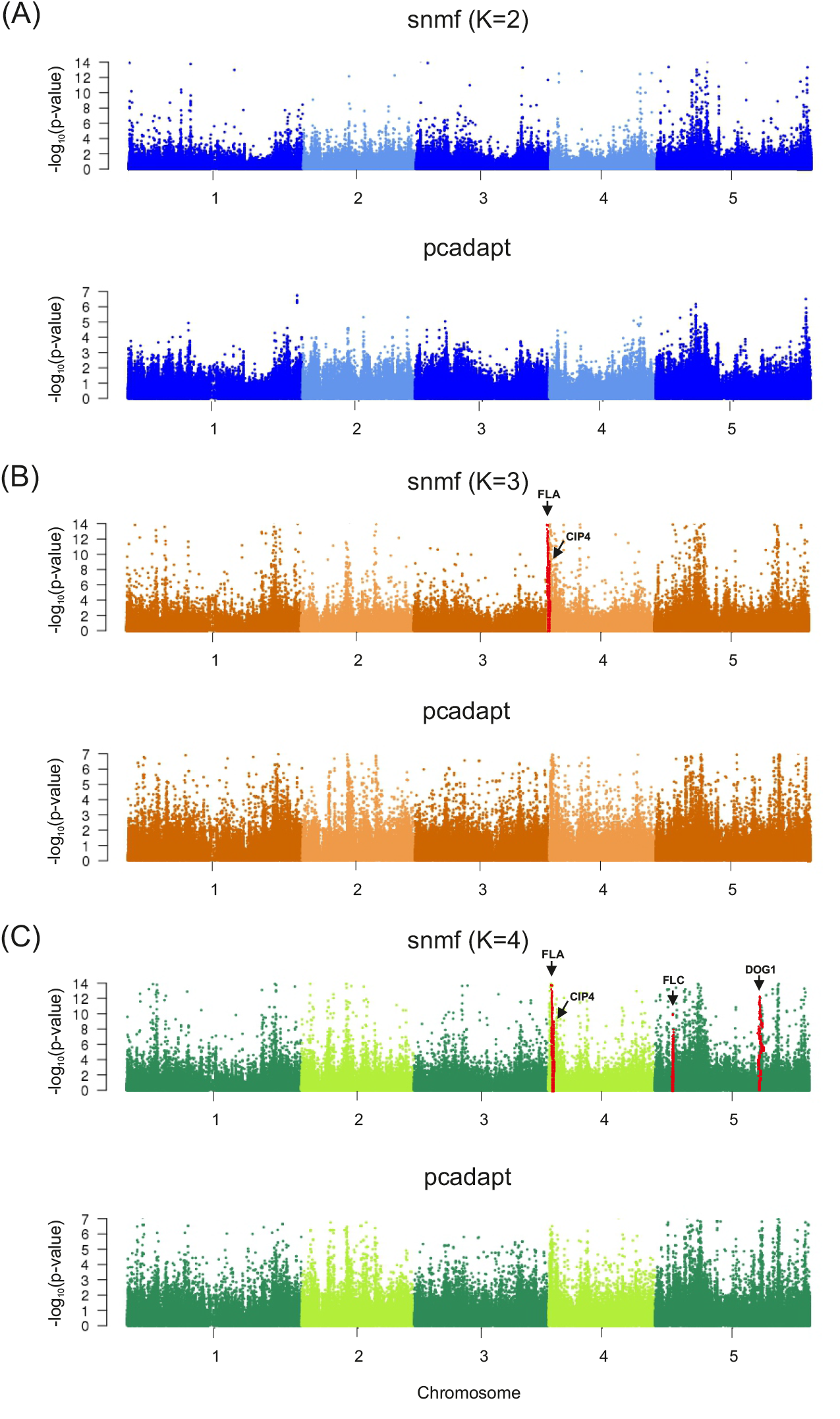
Manhattan plots of minus log10(p-values) for *A. thaliana*. Tests using (A) `snmf` with *K* = 2 ancestral populations and `pcadapt` with 1 principal component, (B) `snmf` with *K* = 3 ancestral populations and `pcadapt` with 2 principal components, (C) `snmf` with *K* = 4 ancestral populations and `pcadapt` with 3 principal components.

